# Genomic insights and breeding strategies for nixtamalization moisture content in hybrid maize

**DOI:** 10.1101/2025.04.23.650239

**Authors:** Michael J. Burns, Sydney P. Berry, Molly Loftus, Amanda M. Gilbert, Candice N. Hirsch

## Abstract

**CORE IDEAS:**

- Nixtamalization moisture content can be selected early in breeding programs using NIR spectroscopy.
- Yield does not significantly correlate with nixtamalization moisture content in diverse or elite populations.
- Additive and dominance gene action impact nixtamalization moisture content in hybrid maize.
- Genomic prediction can be used to assess nixtamalization moisture content early in hybrid maize breeding.

Nixtamalization moisture content, a measure of the quantity of water absorbed during the nixtamalization of a grain such as maize, has a large impact on the end-quality of masa-based products. An application to predict nixtamalization moisture content from raw inbred and hybrid maize grain was recently developed, but its utility in a breeding context has not been assessed. Important breeding considerations for nixtamalization moisture content were assessed in diverse maize hybrids, modern commercial hybrids, and historically high-acreage hybrids grown in up to three environments across two years. This study demonstrated that nixtamalization moisture content is heavily influenced by growing conditions, but sufficient genetic variance is present to allow breeders to make gains from selection. Contrary to prior theory, there was no substantial correlation between nixtamalization moisture content and yield suggesting breeders can select for both traits without negatively impacting either trait. Both additive and dominant genetic action was observed and genomic prediction was able to predict nixtamalization moisture content in hybrids with an average Spearman’s rank correlation coefficient greater than 0.441 and a root mean square error below 0.006. The findings here suggest that nixtamalization moisture content can be selected for early in breeding cycles, allowing breeders to develop improved food-grade maize germplasm without negatively impacting important traits such as yield.

**PLAIN LANGUAGE SUMMARY:** Plant breeders need to understand the biological mechanisms underlying a trait of interest to maximize the efficiency of their efforts. Nixtamalization moisture content is a highly complex trait that is determined by both genetic and environmental factors. In this study, nixtamalization moisture content was assessed in a diverse set of hybrid and inbred maize to understand the biological mechanisms underlying nixtamalization moisture content. The relationship between nixtamalization moisture content and yield was assessed, the genetic architecture and mode of gene action underlying nixtamalization moisture content were evaluated, and the efficacy of genomic prediction in assessing nixtamalization moisture content was determined. The findings of this study will allow breeders to create optimized breeding strategies for nixtamalization moisture content, thus improving the raw materials that are used to produce globally consumed products such as tortillas and tortilla chips.

## 1 INTRODUCTION

Maize is one of the most widely cultivated cereal crops globally, serving as a primary source of food, feed, and industrial raw materials (Bhatnagar et al., 2004; Devi et al., 2024). It plays a crucial role in global food security, providing sustenance for billions of people through direct consumption and indirectly through livestock feed and processed food ingredients (Rooney & Serna-Saldivar, 2015; Serna-Saldivar & Chuck-Hernandez, 2019). One way humans directly consume maize is through masa-based products such as tortillas and tortilla chips (Acosta-Estrada et al., 2023; Molina et al., 2023). These products are created through a high-temperature, high-pH cooking process called nixtamalization which softens the endosperm through water absorption and removes the pericarp through base hydrolysis (Santiago-Ramos et al., 2018; Sefa-Dedeh et al., 2004). The products created from masa are evaluated on several quality parameters, such as taste, texture, nutritional profile, and machinability (Holmes et al., 2019; Yglesias et al., 2005). The quantity of water absorbed during nixtamalization can impact each of these quality parameters (Holmes et al., 2019; Sahai et al., 2001). Nixtamalization moisture content is a complex trait that is largely driven by the composition of the raw kernels when cooking parameters are held constant (Burns et al., 2021), which can be influenced by both genetics and environmental effects (Flint-Garcia et al., 2009; Renk et al., 2021).

Breeding for quality traits can be difficult due to their relationship with other traits of interest such as yield. For example, protein content is known to have a negative correlation with yield, and starch content is known to have a positive correlation with yield across a number of species (Dorsey-Redding et al., 1991; Fukai & Mitchell, 2024; Geyer et al., 2022; Lloyd & Kossmann, 2018; Simmonds, 1995). It is known that starch granules swell to absorb water during nixtamalization, and that protein bodies restrict starch granule swelling (Gutiérrez-Cortez et al., 2022; Holmes et al., 2019; Ratnayake & Jackson, 2006; Santiago-Ramos et al., 2018). By extension, it is hypothesized that increasing yield would increase nixtamalization moisture content, and decreasing yield would decrease nixtamalization moisture content. The relationship between nixtamalization moisture content and yield has not been studied previously, but other species have successfully been bred for increasing yield while maintaining quality traits including baking quality in wheat (Fradgley et al., 2023; Michel et al., 2018; Souza et al., 2002), fatty acid profiles in soybean (Clemente & Cahoon, 2009; Fehr, 2007; Lee et al., 2007), amino acid profiles in pulses (Shahzad et al., 2021; Smartt et al., 1975) and nutritive value in forages (Shenk, 1977). Breeding techniques such as index selection allow breeders to consider multiple traits, even negatively correlated traits, when making selections to improve both quality and agronomic traits (Bernardo, 2014; Yusuf et al., 2024).

To improve nixtamalization moisture content and related masa-quality traits in breeding germplasm, high-quality phenotyping is needed on scales that fit early cycles of breeding programs where sample sizes are small and the number of samples is large (Bernardo, 2014; Schoemaker et al., 2024). However, breeding companies currently make selections late in the breeding process using proxy tests such as kernel hardness, density, or dent depth as these traits tend to correspond to cooking quality (Burns et al., 2021; Holmes et al., 2019; Serna-Saldivar et al., 1993). Eventually, a very limited number of lines that are considered for commercialization are tested in pilot plants that require 500 to 1,000 pounds of grain per sample (Burns et al., 2021). Most of the lines that are sent to manufacturers for pilot testing fail, and a common reason for failure is the quantity of moisture that is absorbed during nixtamalization. An application called CHIP-NMC was recently developed that scales to the needs of breeding programs and is capable of providing accurate estimates of nixtamalization moisture content in diverse maize populations in a high-throughput manner based on near-infrared (NIR) spectra (Burns et al., 2025).

Before breeding for nixtamalization moisture content, it is helpful to understand the genetic basis for the trait including understanding the mode of gene action and the genetic architecture. The genetic basis of traits can heavily influence mating schemes and selection strategies of a breeding program (Gao et al., 2023; Tang et al., 2024; Yi et al., 2019). For example, analyses of fruit quality in apples (Kumar et al., 2015) and grain quality in rice (Ren et al., 2023) have shown additive gene action explains up to 30% of variation in quality traits, and dominance gene action explains only 16% of variation. The mode of gene action for nixtamalization moisture content has not been previously studied and the genetic architecture has only been studied in an inbred context (Burns et al., 2021). Given that maize is a hybrid species in production settings, it is possible that both additive and dominance modes of gene action could affect nixtamalization moisture content in commercial germplasm. Utilizing hybrids when studying the genetic basis of nixtamalization moisture content would allow the dominance genetic architecture to be studied as well. A more thorough understanding of the genetic architecture of nixtamalization moisture content can provide breeders with markers to monitor throughout the early stages of selection (Gao et al., 2023; Jighly, 2024; H. Wang et al., 2017).

In addition to identifying genetic loci associated with nixtamalization moisture content, genomic prediction models could allow breeders to make informed selections in even earlier breeding cycles (Schoemaker et al., 2024; F. Wang et al., 2024). Nixtamalization moisture content is a highly quantitative trait based in part on genes related to grain composition (Gutiérrez-Cortez et al., 2022; Sefa-Dedeh et al., 2004). This complexity matches the assumptions of many genomic prediction models that follow the infinitesimal model (Barton et al., 2017). However, due to dependence of nixtamalization moisture content on composition (Burns et al., 2025; Sefa-Dedeh et al., 2004), it is possible that genomic prediction accuracy could be reduced by environmental effects seen in grain composition (DeSalvio et al., 2024; Renk et al., 2021). While no studies have previously assessed the ability to genomically predict nixtamalization moisture content in maize, success has been observed for quality traits in other species including baking quality in wheat (Fradgley et al., 2023; Michel et al., 2018), fatty acid profiles in soybean (F. Wang et al., 2024), and fruit quality in apple (Kumar et al., 2015).

This study looks at genetic mechanisms and breeding opportunities for nixtamalization moisture content in hybrid maize. To this end, the variance of nixtamalization moisture content in three hybrid panels was partitioned to quantify the effect genetics, environment, and the interaction has on nixtamalization moisture content. The relationship between nixtamalization moisture content and yield was assessed to quantify the expected change in either trait when selecting for the other. The genetic mode of action influencing nixtamalization moisture content was studied through parent-hybrid comparisons and a genome-wide association study. Lastly, a proof of concept genomic prediction model was developed and validated for nixtamalization moisture content in hybrid maize to determine the efficacy of this approach for use in breeding programs. This study applies modern breeding techniques to nixtamalization moisture content and demonstrates that nixtamalization moisture content can be genetically improved without negatively impacting desirable agronomic traits such as yield.

## 2 MATERIALS AND METHODS

### 2.1 Plant Materials and Growth Conditions

The Wisconsin Diversity Panel hybrids, their parent inbred lines, and the panel of commercial hybrids that were used in the development of CHIP-NMC (Burns et al., 2025), were utilized for the analyses in this study. Briefly, a total of 560 diverse hybrids, generated using ∼280 inbred lines from the Wisconsin Diversity Panel (Hansey et al., 2011) as female parents, were crossed with both B73 and Mo17 as the pollen donor. The crossed hybrids, as well as the inbred parent lines, were planted at the same time in Summer 2022 and 2023 at the Agricultural Experiment Station in St. Paul, MN (Burns et al., 2025). Both the hybrid and inbred experiments were designed to include two replicates per year in a randomized complete block design where genotypes were blocked by early and late flowering time within replicates. Total grain was collected from ten consecutive plants in the middle of each plot and dried in a forced air oven at 63°C for seven days. The commercial hybrids were also grown in 2022 and 2023 in St. Paul, MN, and Waseca, MN. The panel of 10 commercial hybrid lines were grown in a randomized complete block design with two replicates in each location in each year. Each plot was planted with four rows, and grain subsamples were collected from the middle two rows after harvest with a plot combine (Supplemental Table S1).

Additionally, a panel of 15 hybrids with historically high acreage in the Midwest, United States, from 1950 to 2020 was assessed (Supplemental Table S1). The historical hybrids were grown in a randomized complete block design with three planting dates that each included two replicates in each location within each year. The experiment was planted in two-row plots on May 11, May 19, and May 31 in St. Paul in 2022, May 24, June 3, and June 10 in Waseca in 2022, May 17, May 26, and June 5 in St. Paul in 2023, and May 22, June 1, and June 8 in Waseca in 2023. Grain subsamples were collected after harvest with a plot combine. Grain samples were harvested on November 22 in St. Paul in 2022, October 27 in Waseca in 2022, November 4 in St. Paul in 2023, and November 9 in Waseca in 2023. A subsample of retained grain was dried at 63°C in a forced air drier for seven days.

### 2.2 Predicting nixtamalization moisture content from NIR spectroscopy

Nixtamalization moisture content was predicted by uploading near-infrared (NIR) spectral data into CHIP-NMC (Burns et al., 2025). The NIR spectra of the Wisconsin Diversity Panel hybrids and inbreds, and commercial hybrids were previously collected (Burns et al., 2025). NIR spectra for the historical hybrids were collected following the method of Burns et al. (2025). Briefly, samples were finely ground using a Perkin Elmer LM3610 grinder and a subset of the ground sample was scanned with a Perten DA7250 spectrometer on the small cup (∼30g), rotating setting. The spectra were output as every fifth waveband from 950nm to 1650nm (Supplemental Table S1). The NIR spectra were uploaded into CHIP-NMC (Burns et al., 2025), with the hybrid model selected for the hybrid plots, and the inbred model selected for the inbred plots. The minimum reported value was set to zero, and the maximum reported value was set to one for unrestricted sample inclusion beyond extrapolation removal. Predictions beyond the training set of the hybrid or inbred model were not included in the output. Nixtamalization moisture content predictions from CHIP-NMC were downloaded and combined with the associated spectra in R v4.0.3 (R Core Team, 2022).

### 2.3 Calculating proportion of variance explained by experimental factors

The distribution of nixtamalization moisture content predictions were plotted out for each population in each year. The proportion of variance explained by experimental factors was calculated with the lme4 package v1.1-26 (Bates et al., 2015) using equation 1 for the Wisconsin Diversity Panel hybrids and equation 2 for the commercial and historical hybrids. For both equations, y is the trait value, μ is the mean value, g is the random genotype effect, e is the random year effect, g x e is the random genotype-by-year interaction, e/r/b is the random block within rep within year effect, and ε is the residual error. Because the commercial and historical hybrid panels were grown at multiple locations in both years, equation 2 additionally includes l as the random location effect, g x l as the random genotype-by-location effect, and e/l/r as the random rep within location within year effect.

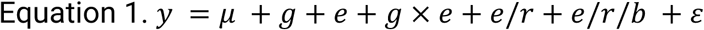

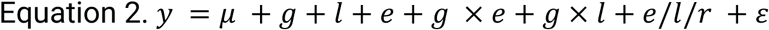

Due to the difference in nixtamalization moisture content between years, and the amount of variation explained by year, the proportion of variation explained by experimental factors was also calculated separately for each year using equation 3 for the Wisconsin Diversity Panel hybrids and equation 4 for the commercial hybrid and historical hybrid panels. Variables for mean value (μ) and random effects from genotype (g) were maintained from equations 1 and 2. In equation 3, r is the random block effect and r/b is random block within rep effect. In equation 4, l/r is random rep within location effect. Subsequent analyses were also performed separately for each year.

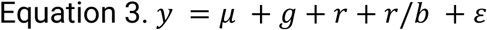

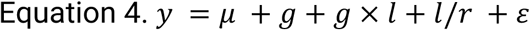

### 2.3 Relationship between nixtamalization moisture content and yield

Yield was determined for each population during harvest by mechanical shelling for the Wisconsin Diversity Panel hybrids and inbreds and with a plot combine for the commercial and historical hybrids. For mechanical shelling, all grain-bearing ears from ten sequential representative plants within a plot were harvested and mechanically shelled. Grain mass, test weight, moisture, and stand count information were used to extrapolate the plot yield for plots with at least 75% germination. After the grain quantity per plot was determined and corrected for moisture, it was extrapolated from a per-plot basis to a per-hectare basis for comparison in standard units (tons/hectare). Due to the size of the Wisconsin Diversity Panel experiments, the yield of each plot was adjusted according to the average difference of checks in its block-rep combination to the overall check average. For plot combine shelling, plots were harvested with a plot combine that provided moisture-corrected yield in bushels per acre. The yield was converted to standard units (tons/hectare) for comparison. The commercial and historical hybrid experiments were spatially restricted and therefore did not include spatial correction with checks.

To determine the genotypic relationship between nixtamalization moisture content and yield, best linear unbiased estimates (BLUEs) were calculated in each population within each year using equation 3 for the Wisconsin Diversity Panel hybrids and equation 4 for the commercial hybrids and historical hybrids. For all experiments, genotype was a fixed effect and all other factors were random. The lme4 package v1.1-26 (Bates et al., 2015) was used to model both nixtamalization moisture content and yield and BLUEs were extracted as the genotype value plus the model intercept in each model. The Pearson’s correlation coefficient between nixtamalization moisture content BLUEs and yield BLUEs was determined for each population in each year.

### 2.4 Impact of inbred parents on hybrid nixtamalization moisture content

The impact of inbred parents on hybrid nixtamalization moisture content was assessed using the Wisconsin Diversity Panel inbreds and hybrids. This comparison was made using the nixtamalization moisture content BLUEs for the hybrids as described above, and by calculating the nixtamalization moisture content BLUEs for the inbred parents using equation 3, where genotype was treated as a fixed effect and all other factors were random effects. After the BLUEs for both populations were calculated, the inbred BLUEs for each hybrid cross were used to determine the high-parent values and calculate the mid-parent values. The deviation of each hybrid relative to the high-parent and mid-parent value was plotted in a density plot and t-tests were performed to determine if the hybrids deviated significantly (p < 0.05) from the high-parent or mid-parent values. In addition to assessing the deviation from high-parent and mid-parent values, the BLUEs of inbreds and hybrids were plotted in a reaction norm plot to assess the presence of additive and non-additive effects that inbred combinations might have.

To determine if hybrid offspring could be directly predicted from the phenotype of their inbred parents, a linear regression model was trained on 80% of the hybrid genotypes using the lm function in R v4.0.3 (R Core Team, 2022) using equation 5. The model was iterated 100 times to account for partitioning variation affecting the accuracy of predicting hybrid nixtamalization moisture content BLUEs from inbred nixtamalization moisture content BLUEs within each year. The coefficients of each parent and the interaction term were used to determine if additive and dominant modes of action were present.

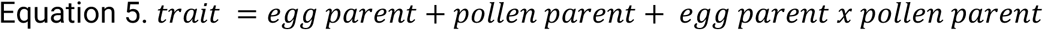

### 2.5 Genetic architecture of nixtamalization moisture content in hybrid maize

The genetic architecture of nixtamalization moisture content in a hybrid system was determined by performing a genome-wide association study on the Wisconsin Diversity Panel hybrid’s nixtamalization moisture content BLUEs. TASSEL 5 (Bradbury et al., 2007) was used through the command line interface to convert the VCF file of previously published SNP data for the Wisconsin Diversity Panel (Qiu et al., 2021) into hapmap format. Heterozygous markers were set to missing (‘NN’) as it is unclear which allele would be passed on to hybrid progeny. The marker data of the inbred lines was then used to computationally derive the hybrid genotypes based on the inbred parent pedigree by concatenating the single unique marker for both parents at every locus. The markers were filtered for a minor allele frequency greater than 5%, and missing data less than 15%. The hapmap file was then numericalized with either additive or dominance coding using GAPIT v3.5 (Lipka et al., 2012). The additive coding returned a value of zero or two for homozygous markers and one for heterozygous markers while the dominance coding returned a value of 0 for homozygous markers and one for heterozygous markers. The additive and dominance coded markers were used to determine the percent phenotypic variance explained by marker data. The data was filtered to remove any sites with missing alleles and converted to either an additive or dominance genetic relatedness matrix following previous work (Burns et al., 2021). The additive and dominance genetic relatedness matrix and phenotypic BLUEs were input into GCTA v1.94.0 (Yang et al., 2011) to determine the proportion of phenotypic variance explained by additive or dominance genetics.

To determine the population structure of the Wisconsin Diversity Panel hybrids, a principal component analysis was performed on the original hapmap using GAPIT v3.5 (Lipka et al., 2012). Following this, the phenotype file of nixtamalization moisture content BLUEs for each year, the numericalized hapmap, the genetic map, and the population structure files were filtered to ensure the contents were the same and in the same order. The GWAS methods followed previous work on inbred grain composition and nixtamalization moisture content (Burns et al., 2021; Renk et al., 2021). Briefly, FarmCPU v1.02 (Liu et al., 2016) was used to perform GWAS on additive and dominant coded hapmap files to find marker associations in each year, which provided the significance and marker effect size of each marker. The coordinates of significant markers were used to find the closest gene upstream or downstream of the marker in the MaizeGDB genome browser (Woodhouse et al., 2021). The gene sequence, exon sequence, and amino acid sequence from the annotation in MaizeGDB (Woodhouse et al., 2021) were used to determine the putative functional annotation using NCBI’s standard non-redundant (nr) sequence query database (BLASTN = gene sequence, BLASTX = exon sequence, and BLASTP = amino acid sequence; Altschul et al., 1990). The most common annotation was recorded for each BLAST query.

### 2.6 Genomic prediction model

A genomic prediction model was developed using the results of the GWAS described above. Markers were selected based on their association with nixtamalization moisture content. The twenty-five most significant markers from each chromosome were iteratively selected within each year and from both the additive and dominance GWAS models. To reduce redundancy within year and model type, after the most significant marker was selected, markers within 500Kb of the marker were removed from consideration. This threshold was determined following a previously described method (Della Coletta et al., 2023). Briefly, the background LD and LD decay rate for the Wisconsin Diversity Panel was calculated by determining the LD between 50 random markers across all chromosomes with PLINK v1.90b6.10 (Purcell et al., 2007) a. The window in which the median LD fell below the background LD was used as the redundancy threshold between markers.

After the hapmap dataset was reduced according to GWAS significance, heterozygous sites were converted to missing values (NN) as it is unknown which allele is present in the hybrid. Imputation was then performed on all missing data, including once heterozygous sites, through a decision tree model which was trained to predict missing data for each site based on the allelic states of 250 genotypes that did not have missing data. The fully imputed dataset was used to computationally derive the genotypes of the hybrid offspring by concatenating the unique marker values of the inbred parents at each site. The hybrid hapmap was numericalized where the first allele listed in the hapmap allele column was converted to zero, the second allele listed in the hapmap allele column was converted to two, and the heterozygous state was converted to one (Supplemental Table S2). A pairwise filtering was performed between the numerical hapmap genotype and phenotype files to ensure prediction models were built using only data that had both genetic and phenotypic information for 2022 and 2023. This filtering resulted in 443 hybrids included in the analysis for both years (Supplemental Table S2).

A ridge regression best linear unbiased prediction (rrBLUP) model was created through the RRBLUP R package v4.6.1 (Endelman, 2011) for nixtamalization moisture content BLUEs. The model was tested by splitting the dataset into 10 folds for each year that were used to cross-validate. The rrBLUP model was then trained on 9 folds and used to predict the remaining fold for the respective year. This was repeated until each fold was used as the performance testing set. This cross-validation scheme was replicated 10 times to improve the prediction accuracy assessment. The rrBLUP models were trained to predict nixtamalization moisture content BLUEs as a function of the genetic information in a CV1 design where unknown genotypes were predicted in a known environment. Spearman’s rank correlation coefficient and root mean squared error were collected for each year and recurring parent combination. The average prediction of each hybrid in each year was determined and plotted to illustrate differences between the prediction performance of each recurring parent in each year.

## 3 RESULTS AND DISCUSSION

### 3.1 Nixtamalization moisture content varies significantly across years

The hybrid model in CHIP-NMC (Burns et al., 2025) was used to predict the nixtamalization moisture content from NIR spectral data in the Wisconsin Diversity Panel hybrid experiment, the commercial hybrid experiment, and the historical hybrid experiment, and used to determine how variation partitioned across years and experimental locations. These predictions were roughly normally distributed across and within each year, with overall means of 0.427 (0.377 to 0.467) for Wisconsin Diversity Panel hybrid experiment, 0.426 (0.400 to 0.456) for the commercial hybrid experiment, and 0.435 (0.411 to 0.456) for the historical hybrid experiment (Figure 1A). In each population, 2022 samples had a higher average nixtamalization moisture content than 2023 by an average of 0.0116 (0.0103 to 0.0127; Figure 1A). This magnitude of difference across years was consistent with previous findings (Burns et al., 2021), demonstrating that growing environment has a large impact on the nixtamalization moisture content of grain. These differences are likely due to shifts in grain composition, the distribution of macromolecular components throughout the kernel, and kernel morphology, but more research is needed to parse the causative environmental factors and associated compositional changes that lead to altered maize nixtamalization moisture content.

**Figure 1.**
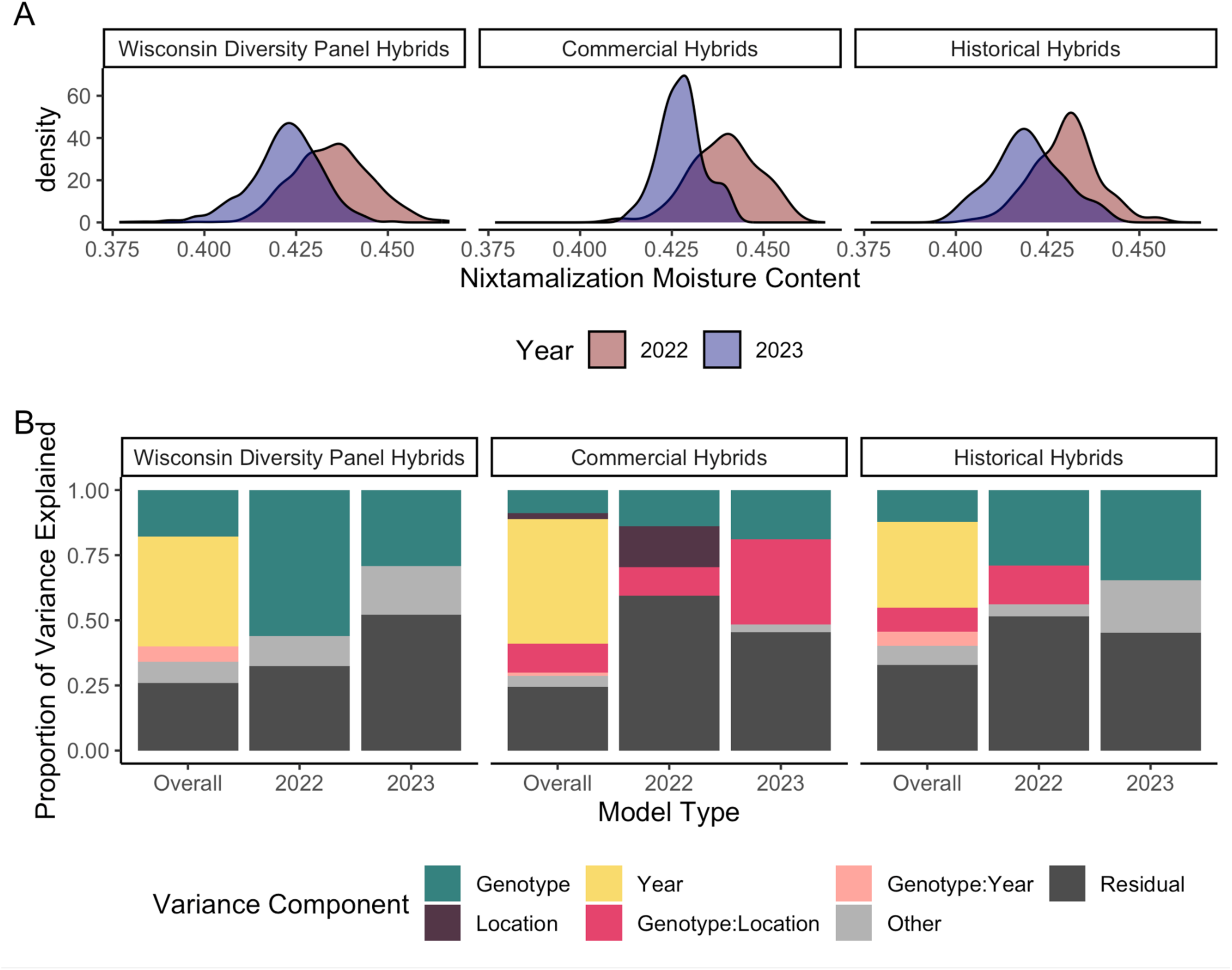
Distribution and variance partitioning of nixtamalization moisture content. (A) Density distribution of nixtamalization moisture content in the Wisconsin Diversity Panel hybrids, commercial hybrid panel, and historical hybrid panel for both years of the study. (B) Proportion of variance explained by each experimental factor determined by mixed linear modeling. The ‘other’ category includes rep within year, rep within location within year, location by year, and block within rep within year effects.

The proportion of variance explained by experimental factors was also determined for each population to further quantify how much environment and genotype impacted nixtamalization moisture content in each population. The results of the mixed linear model analysis showed that year explained the largest proportion of variance when both years were included in the mixed linear model (Figure 1B). When both years were analyzed separately, genotype became the largest source of non-residual variation (Figure 1B). The variance parameters were used to determine heritability of nixtamalization moisture content in each population. The entry mean heritability in the Wisconsin Diversity Panel hybrid experiment was 0.655, in the commercial hybrid experiment it was 0.440, and in the historical hybrid experiment it was 0.682 when years were combined. The heritabilities observed in this study are lower than what was observed in previous research on nixtamalization moisture content conducted in inbred maize germplasm (Burns et al., 2021), but similar to heritabilities for quality traits in other species like apple (Kumar et al., 2015) and wheat (Souza et al., 2002). These results demonstrate that genetic variation exists in hybrid maize germplasm for nixtamalization moisture content, and suggest that breeders should be able to make gains from selection for nixtamalization moisture content.

### 3.2 Yield and nixtamalization moisture content are not correlated

Yield is one of the most important traits that breeders select for in maize, and it is known to be significantly correlated with traits such as starch and protein content (Dorsey-Redding et al., 1991; B. Ma et al., 2023; Renk et al., 2021), which can influence nixtamalization moisture content (Burns et al., 2021, 2025; Santiago-Ramos et al., 2018; Sefa-Dedeh et al., 2004). To determine the relationship between yield and nixtamalization moisture content, the Pearson’s correlation coefficient was calculated between BLUEs for yield and BLUEs for nixtamalization moisture content. A significant correlation was observed in the Wisconsin Diversity Panel in 2023, but the Pearson correlation coefficient was only 0.1 (Figure 2). The minimal correlation in the Wisconsin Diversity Panel hybrid experiment could be due to the fact that it is a diverse panel that has not undergone selection for yield. The relationship between nixtamalization moisture content and yield was also evaluated in the commercial hybrid experiment and historical hybrid experiment, both of which contain hybrids that have undergone significant selection for yield improvement. In both experiments no significant correlation between yield and nixtamalization moisture content was observed in either year (Figure 2). While the commercial hybrids had an overall higher yield than the Wisconsin Diversity Panel, the range of yield values was much lower (∼3 tons/hectare), which could reduce the ability to observe a statistical relationship between nixtamalization moisture content and yield. The historical hybrid experiment, however, included hybrids with considerably more range in yield (∼5 tons/hectare), and still no significant correlation between nixtamalization moisture content and yield was observed (Figure 2). The lack of significant correlations suggests that yield and nixtamalization moisture content are not necessarily synergistic or antagonistic, and that breeders can continue to breed for improved nixtamalization moisture content without detriment to yield.

**Figure 2.**
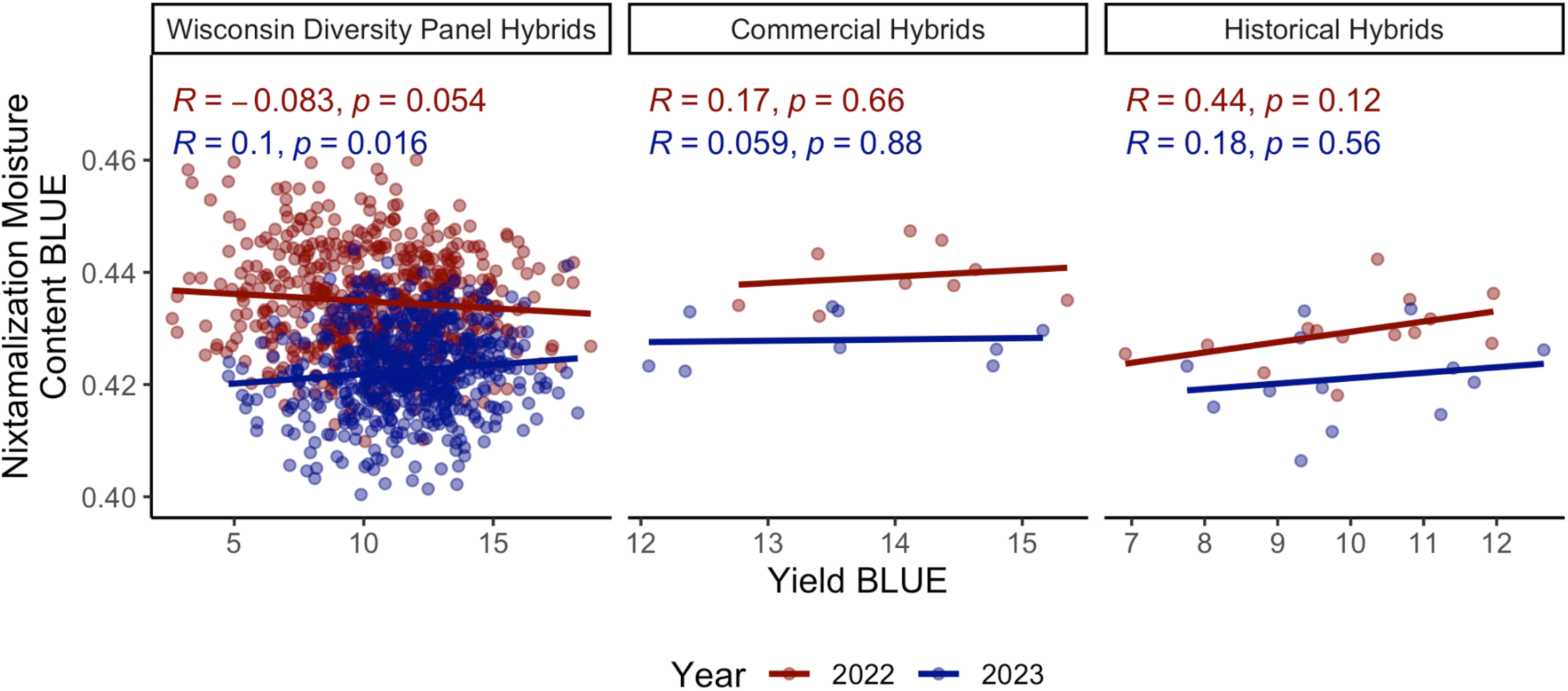
Correlation of yield and nixtamalization moisture content. The correlation of yield BLUEs and nixtamalization moisture content BLUEs for the Wisconsin Diversity Panel hybrids, commercial hybrid panel, and historical hybrid panel. Correlations within each year were calculated with Pearson’s correlation coefficient.

### 3.3 Additive and dominance gene action is observed between inbred parents and hybrid progeny

To assess the effect of genetic additivity and dominance on nixtamalization moisture content, the Wisconsin Diversity Panel hybrids and the inbreds associated with the crosses were compared. Because year explained the largest proportion of variance (Figure 1B), it was controlled for by comparing hybrid and inbred data within respective years. Additivity and dominance effects were first assessed through a reaction-norm plot, which showed that differences in pollen parent performance transferred to hybrid performance, particularly in 2022, suggesting that additive modes of gene action impact nixtamalization moisture content (Figure 3A). It is also notable that hybrids tend to have more extreme nixtamalization moisture content than their inbred parents, suggesting that dominance effects are also occurring (Figure 3A).

**Figure 3.**
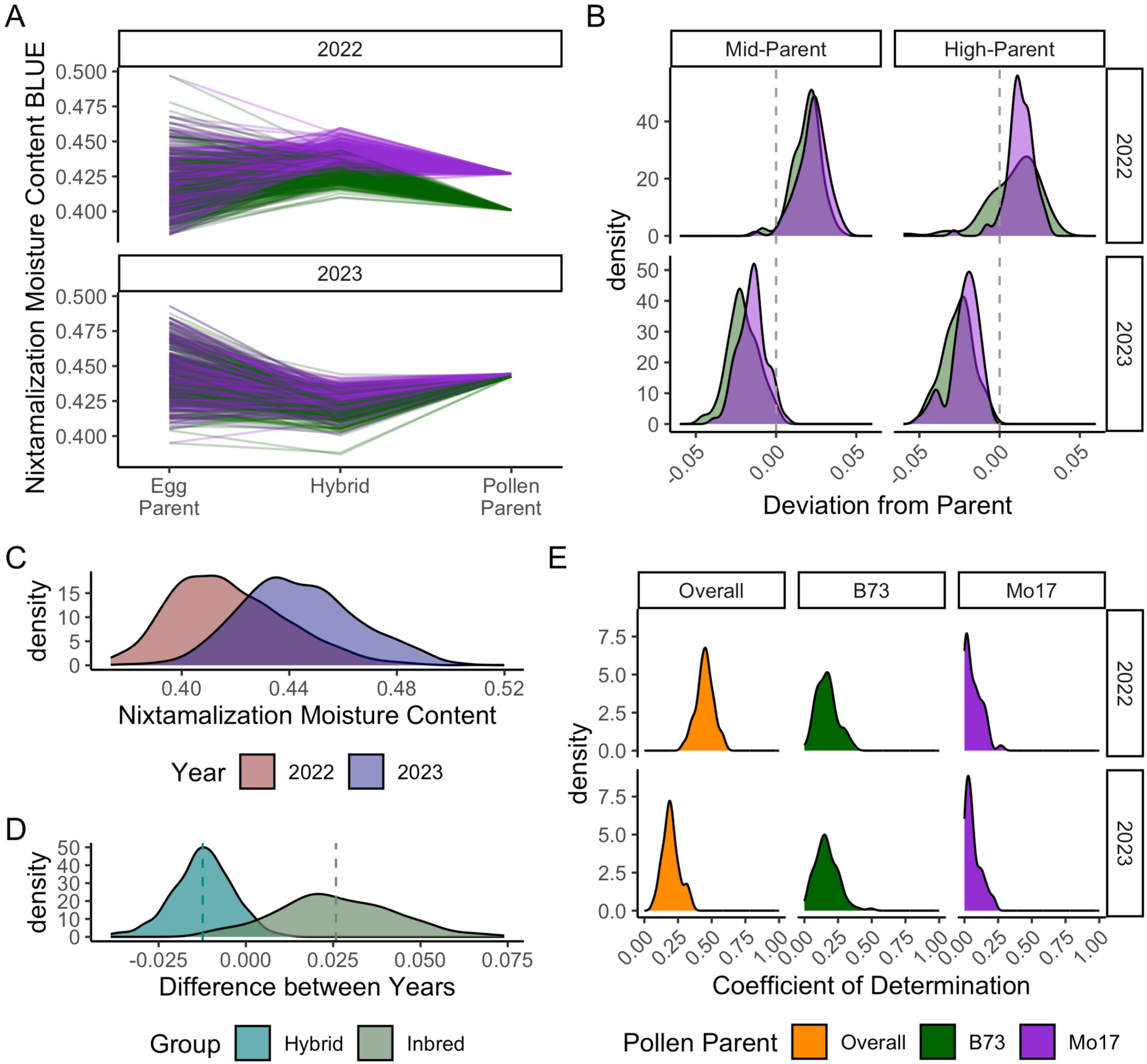
Breeding considerations of nixtamalization moisture content. (A) Reaction norm plots between inbred parents and hybrid progeny. (B) Nixtamalization moisture content BLUEs of the Wisconsin Diversity Panel hybrids were compared to the mid-parent and high-parent nixtamalization moisture content BLUEs of the Wisconsin Diversity Panel inbreds to understand the additivity and dominance gene actions that might be underlying the trait. (C) Distribution of nixtamalization moisture content in the Wisconsin Diversity Panel inbreds in 2022 and 2023 as predicted by CHIP-NMC. (D) Distribution of differences in nixtamalization moisture content across years (2023 - 2022) in inbred and hybrid experiments. Vertical dashed lines represent the average difference for each group.(E) Prediction of nixtamalization moisture content BLUEs of the Wisconsin Diversity Panel hybrids performed through linear modeling based on parental nixtamalization moisture content BLUEs and repeated 100 times to determine prediction performance variability.

To quantify dominance effects, the deviation of the hybrid offspring from the midparent and high-parent value of the inbred parents was evaluated. In 2022, hybrid nixtamalization moisture content was generally higher than both the midparent (mean difference = 0.0194, p-value = 2.31×10^-187^) and high parent values (mean difference = 0.0101, p-value = 1.17×10^-50^). Conversely, in 2023, the nixtamalization moisture content of the hybrids were lower than both the mid parent (mean difference = −0.0211, p-value = 5.61×10^-206^) and high parent values (mean difference = −0.284, p-value = 1.21×10^-226^; Figure 3B). This was driven more by changes in the mean of inbred values between the years than between the means of the hybrid values (Figure 1A and Figure 3C,D). It is well documented that hybrid phenotypes are more stable than inbred phenotypes in extreme environments (Ali et al., 2017; Costa et al., 2024; Li et al., 2021). The two environments used in this study were hotter and drier than average throughout most of their growing season, but differed significantly in precipitation during key developmental stages such as grain filling and dry down (Supplemental Figure S1). It is possible that the growing conditions across years and the response of the inbreds to those conditions are what created the larger and opposite shift in nixtamalization moisture content in the inbreds compared to the hybrids. Future research is needed to understand the full impact of growing conditions on nixtamalization moisture content.

A linear regression model was created to determine the significance of additive and non-additive relationships between inbred parents and their hybrid progeny, and to determine if hybrid nixtamalization moisture content can be predicted based on the BLUEs of both inbred parents. The model was trained with a portion of the data to allow for prediction of the remainder of the data, and this was iterated 10 times. The pollen parent term was significant in 94.5% of model iterations, the egg parent was significant in 82% of model iterations, and the interaction of pollen parent and egg parent was significant in 73% of model iterations. The prediction accuracy, measured by Pearson’s coefficient of determination, was modest when considering both B73 and Mo17 crosses in 2022 (mean = 0.449, range = 0.289 to 0.602), and relatively low in 2023 (mean = 0.198, range = 0.0644 to 0.351; Figure 3E). Models were also trained for each of the pollen parents separately. The model trained and evaluated on B73 crosses performed better than Mo17 models in both 2022 (mean difference = 0.0995) and 2023 (mean difference = 0.102; Figure 3E). Overall, these results indicate that the performance of the inbred parents, even in the same environment, is only slightly predictive of the hybrid performance. Previous research reported that NIR spectroscopy models developed for inbred grain could not be applied to hybrid grain (Burns et al., 2025). It was proposed that differences in kernel composition distribution and morphology could cause the lack of accuracy of the inbred model on the hybrid germplasm (Burns et al., 2025). These differences could also cause the lack of predictive power observed here as the compositional distribution and morphology of hybrid kernels is not expected to be representative of the inbred kernels due to the higher degree of vigor, photosynthetic area, and successful embryo fertilization. These factors can lead to increased quantities of storage molecules such as starch and protein, and tighter packed kernels on an ear, leading to different internal distributions of hard and soft endosperm.

### 3.4 Genetic architecture of nixtamalization moisture content includes additive and dominant markers

A genome-wide association study was performed on the Wisconsin Diversity Panel hybrids after computationally deriving the hybrid genotypes based on the genotypes of the inbred parents for a set of 1.38 million SNPs after all filtering criteria. Both additive and dominance codings of the genotypes were assessed using FarmCPU v1.02 (Liu et al., 2016). Only one significant additive marker and seven significant dominant markers were observed in 2022 (Figure 4; Supplemental Table S3). No significant markers were found in 2023, which could be due to the decreased phenotypic variation between the pollen parents (Figure 3A). While few significant associations were observed, the full set of additive coded SNP data explained 56.3% of the phenotypic variation in 2022 and 36.3% in 2023 and the dominance coded SNP data explained 48.4% in 2022 and 32.2% in 2023. These results collectively indicate that this trait is controlled by many loci of small effect. This is consistent with previous GWAS on compositional traits in maize that may be important for nixtamalization moisture content (Renk et al., 2021), and on the trait of nixtamalization moisture content itself when evaluated in inbred germplasm (Burns et al., 2021).

**Figure 4.**
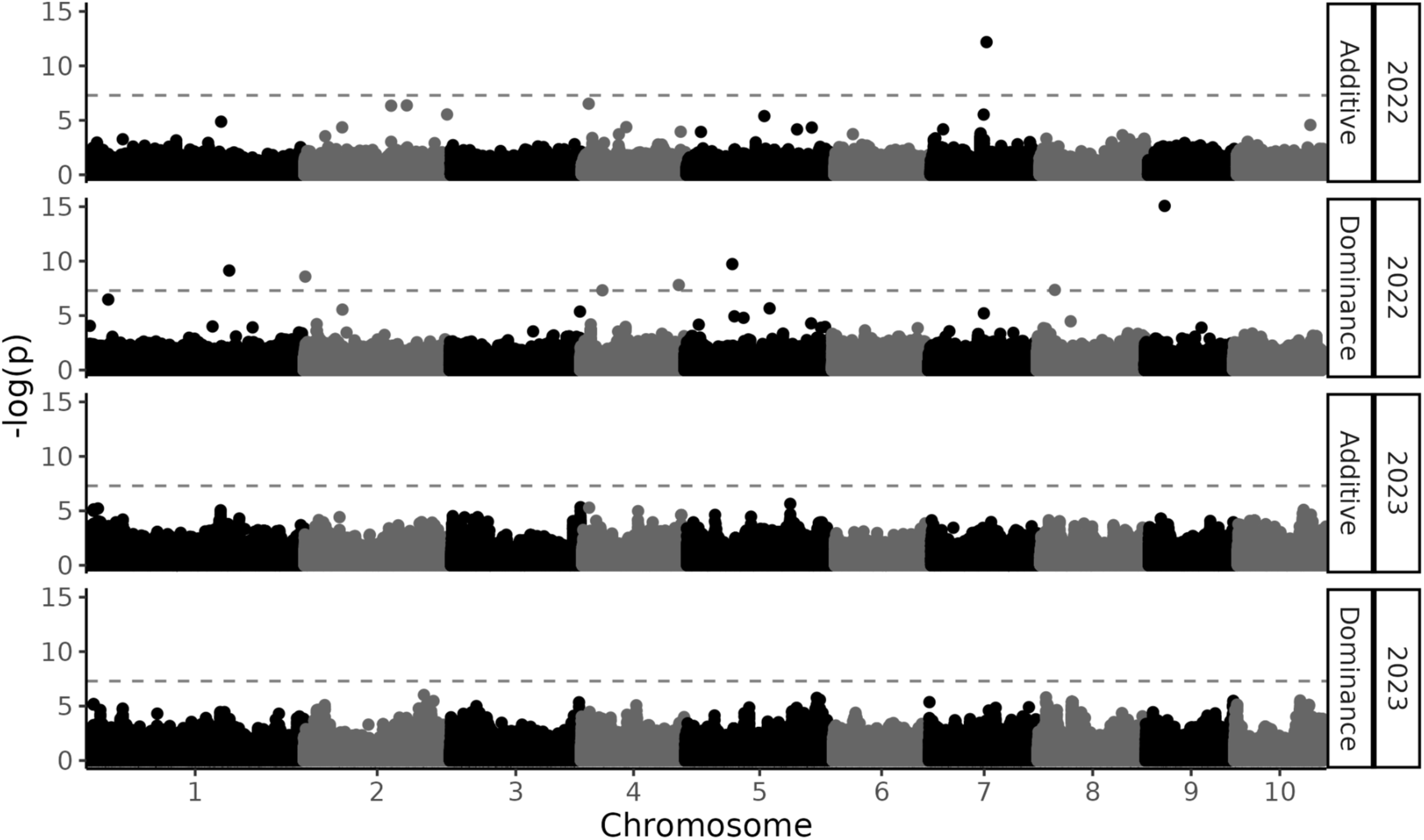
Additive and dominance genetic architecture of nixtamalization moisture content. Genome-wide association studies were performed in GAPIT and FarmCPU to determine the genetic architecture of nixtamalization moisture content in maize. Additive and dominance scores were used to assess both modes of gene action in both years of the study.

Of the significant markers found, four were upstream of the nearest gene, and four were downstream. The average distance to a gene was 23,715 base pairs (2,953 to 54,213; Supplemental Table S3). Six of the genes have putative functional annotations related to environmental stress response through pathways including hormone signaling (Blum et al., 2025; Ren et al., 2023), RNA alternative splicing (J. Ma et al., 2025), and molecular catabolism (Vogel et al., 2008). Although none of the markers or genes reported here overlap those found in a previous study on nixtamalization moisture content in germplasm (Burns et al., 2021), common themes in putative functions did emerge. Environmental stress response was a repeated finding in the previous work as it is in this study. Additionally, there were multiple hormone-signaling pathways observed previously (Burns et al., 2021). These results are consistent with prior work in suggesting that environmental stress response is a major contributor to variation in nixtamalization moisture content that should be considered throughout the breeding process.

### 3.5 Genomic prediction is moderately useful for predicting nixtamalization moisture content

Nixtamalization moisture content is controlled by many small effect loci which is ideal for genomic prediction models, particularly those based on the infinitesimal model. To evaluate the utility of genomic selection for nixtamalization moisture content, genomic prediction models were created for nixtamalization moisture content using the Wisconsin Diversity Panel hybrids. To develop the genomic prediction model, the 1,000 most significant markers with balanced representation across each chromosome were identified. Redundancy between markers was controlled for by calculating background LD (R^2^ = 0.0396) and the base pair window in which the median LD of the population reached this level (500Kb). Of the 1,000 selected markers, 1.7% of the data was heterozygous, and 0.23% of the data was missing.

The 1,000 selected markers were used to train rrBLUP models to predict nixtamalization moisture content BLUEs as a function of the genetic information in a CV1 scheme. Here we refer to CV1 as the prediction of unknown genotypes in known environments. The predicted nixtamalization moisture content values were compared to the observed nixtamalization moisture content BLUEs across and within pollen parent groups for each replicated cross-validation fold (Figure 5A). The correlations across pollen parent groups was moderate to high in both years, averaging 0.792 in 2022 and 0.621 in 2023. Given that pollen parent group differences were noted when comparing hybrid and inbred nixtamalization moisture content (Figure 3A), the performance of genomic prediction within each pollen parent group was also assessed. The within pollen parent group correlations were generally lower within each year than the overall correlation. The average genomic prediction accuracy ranged from 0.441 to 0.623 in each population across 2022 and 2023 (Figure 5A; Supplemental Table S4). This difference in performance for each subgroup compared to the overall performance is due to the difference in nixtamalization moisture content for each pollen parent group, particularly in 2022 (Figure 5B). The average performance of B73 and Mo17 hybrids differed enough that a high correlation could be achieved by accurately predicting the group average. In addition, the combination of a large difference in group average and a moderate dynamic range of predictions in 2022 led to an overall correlation that is not representative of either subpopulation, a phenomenon known as Simpson’s paradox (Sprenger & Weinberger, 2021). It is notable that Mo17 crosses were correctly predicted as having higher nixtamalization moisture content BLUEs than B73 crosses (Figure 5B). These moderate to high correlations between genomically predicted nixtamalization moisture content and NIR predicted nixtamalization moisture content indicate that genomic prediction could allow breeders to select against extremely high or low nixtamalization moisture content in early breeding cycles. The results support using genomic prediction in early stages of breeding when destructive sampling of 30g of grain is not feasible, and incorporating phenotypic selection through NIR spectroscopy in later stages when more grain is available.

**Figure 5.**
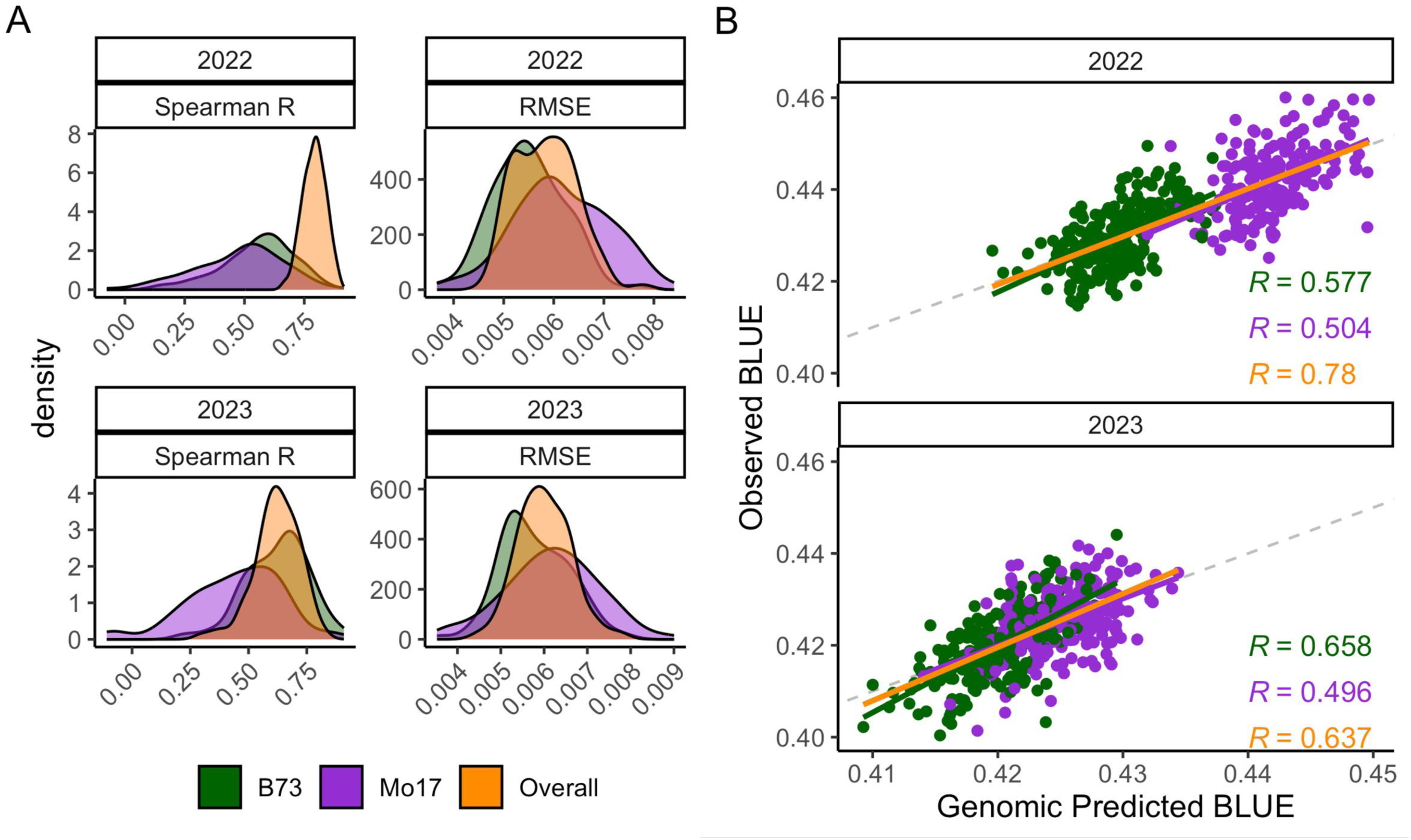
Genomic prediction performance for nixtamalization moisture content. (A) The distribution for each genomic prediction model in a CV1 scheme. The Spearman’s rank correlation coefficient and root mean square error (RMSE) were collected for each pollen parent group as well as overall in each year. (B) The correlation between nixtamalization moisture content and genomic predicted nixtamalization moisture content is based on the average genomic prediction for each genotype over all 10 replications it was predicted in. Gray dashed line represents a 1:1 ratio between predicted and actual trait values.

## 4 CONCLUSION

In this study we assessed the utility of an inbred and hybrid machine learning model on predicting nixtamalization moisture content in realistic hybrid systems. Our results suggest that while environment has a large impact on nixtamalization moisture content, enough genetic variation exists for breeders to select for nixtamalization moisture content. The lack of correlation between yield and nixtamalization moisture content in three different populations suggests that selection pressure on nixtamalization moisture content will not come at the detriment of yield, which is important for both breeders and the producers who grow food-grade varieties. This study also demonstrated the complex relationship between hybrid and inbred parent nixtamalization moisture content, indicating that breeders should be careful when selecting inbred parents based on their nixtamalization moisture content. Breeders should focus their inbred selections on additive modes of gene action and assess hybrids for *per se* phenotypes due to the inconsistent dominance gene action that occurred in this study, which could be due to inbred reactions to the environment. The genome-wide association study performed in this study provided eight markers that highlight the importance of environmental stress response to nixtamalization moisture content, suggesting that breeding for robust stress response could be beneficial for nixtamalization moisture content in hybrids. Finally, genomic predictions had moderate to high correlations across and within subpopulations indicating that breeders can select for nixtamalization moisture content early in plant breeding cycles when the quantity of grain is too limited for NIR spectroscopy, and transition to NIR spectroscopic predictions when the quantity of grain increases in later breeding cycles. The findings of this study display a range of applications and utility for further research into nixtamalization moisture content and provides plant breeders with practical tools that can be incorporated into food-grade maize breeding programs to improve nixtamalization moisture content.

## SUPPLEMENTAL MATERIALS

The data that supports the findings of this study are available in the supplementary material of this article.

## DATA AND CODE AVAILABILITY

Phenotypic and spectroscopic data is available at doi.org/10.1002/cche.10874 (Burns et al. 2025). The raw genotypic data used for GWAS and genomic selection is available at doi.org/10.1101/2020.09.25.314401 (Qiu et al. 2021). The code for this manuscript is available at https://github.com/HirschLabUMN/HNMC_Breeding.

## AUTHOR CONTRIBUTIONS

CNH conceived this experiment. MJB, SPB, ML, AMG conducted the experiments. MJB analyzed the data and visualized the data. MJB and CNH wrote the original draft. All co-authors edited and approved the final manuscript.

## ACKNOWLEDGEMENTS

We thank the Minnesota Supercomputing Institute (MSI) at the University of Minnesota for providing resources that contributed to the research results reported in this paper. This work was funded by PepsiCo, Inc. The views expressed in this manuscript are those of the authors and do not necessarily reflect the position or policy of PepsiCo, Inc. MJB was funded by the University of Minnesota Informatics Institute MnDRIVE Graduate Assistantship and the University of Minnesota Doctoral Dissertation Fellowship.

