## Supplemental Figures for "Genomic insights and breeding strategies for nixtamalization moisture content in hybrid maize"

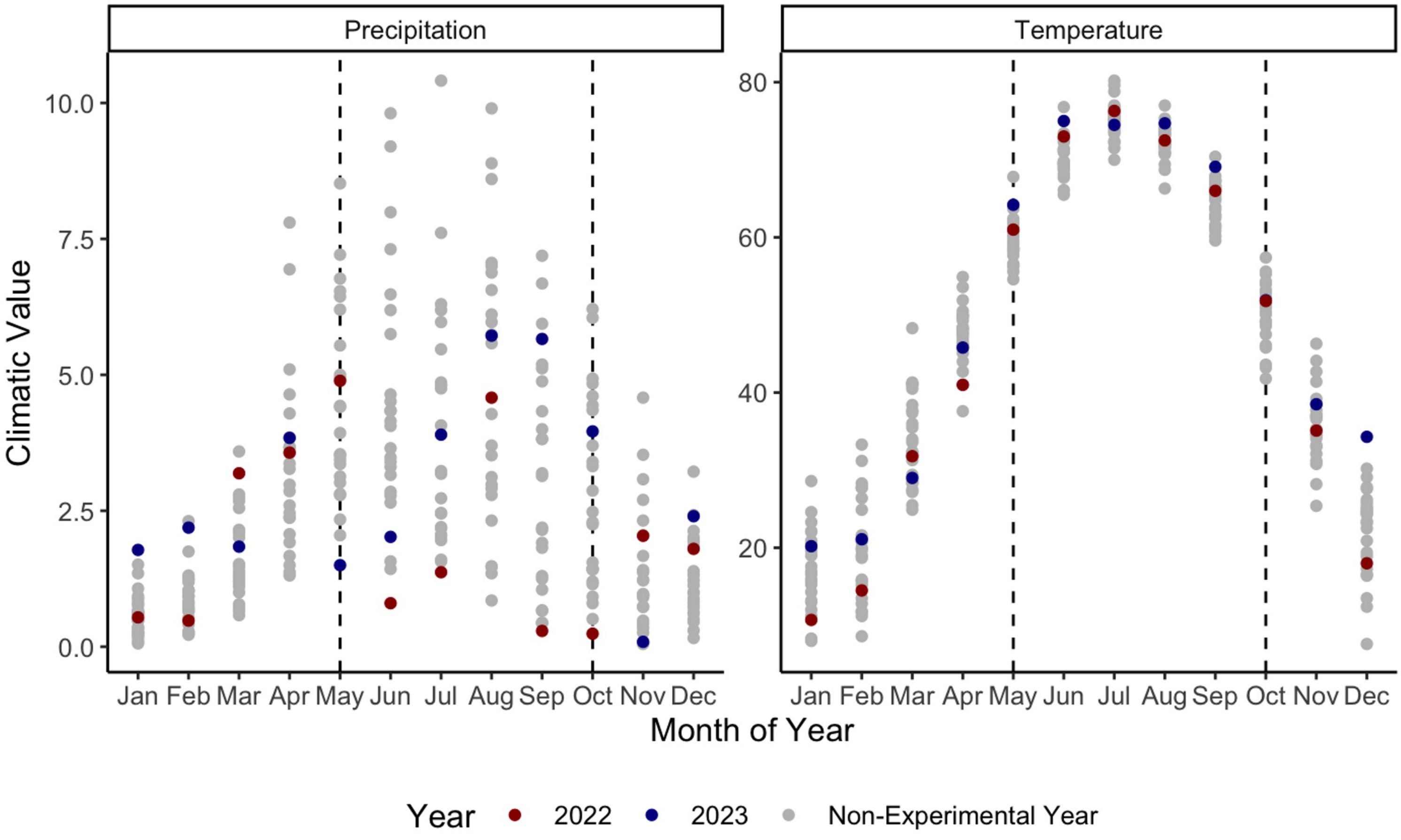


**Supplemental Figure S1. Climatic growing conditions for the Wisconsin Diversity Panel hybrids.** Total monthly precipitation and average monthly temperature for 2000 through 2024 are documented. Vertical dashed lines represent the beginning and end of a typical growing season. Color of points represent the year data was collected, with gray representing years not present in this study.
